# Peucedanol ameliorates LPS-induced inflammation in RAW264.7 cells and CLP-induced sepsis in mice by inhibiting TLR4/myD88/NF-κB pathway

**DOI:** 10.1101/2024.08.02.606445

**Authors:** Qi Yao, Bo-tao Chang

**Author notes:** Corresponding author is to be contacted with, (Qi Yao). Address: 157 Jinbi Road, Xishan District, Kunming 650032, China. They contributed equally to this work.

## Abstract

**Background:** Previously, it has reported that Peucedanol (PEU) possesses anti-bacterial activity. However, its effect and mechanism against inflammation remains unclear.

**Methods:** Isothermal titration calorimetry (ITC) was used to assess binding affinities of PEU to pathogen associated molecular patterns (PAMPs) Kdo2-Lipid A (KLA), oligodeoxynucleotide 1826 (ODN 1826), and peptidoglycan (PGN). A lipopolysaccharide (LPS)-induced RAW264.7 cell inflammation model and a cecum ligation and a puncture (CLP)-induced mouse sepsis model were used to assess efficacy and mechanism of PEU *in vitro* and *in vivo*. 16S ribosomal RNA (16S rRNA) sequencing was used to assay characteristics of intestinal flora of the sepsis mice.

**Results:** PEU had a moderate binding to KLA and ODN 1826. PEU significantly reduced supernatant tumor necrosis factor α (TNF-α) and interleukin 6 (IL-6), and downregulated protein expressions of toll-like receptor 4 (TLR4), myeloid differentiation primary response gene 8 (MyD88), and nuclear factor kappa-B (NF-κB) in the LPS-treated cells. PEU remarkably increased the survival rate, reduced the serum TNF-α and IL-6 levels, attenuated the CLP-induced pathological damage of intestine, increased proliferation-related proteins Bmi1 and Lgr5. Further, the anti-inflammatory effects of PEU were not significantly abolished in the present of chloroquine (CQ). Meanwhile, PEU significantly increased Chao1 index of the intestinal flora at the early stage of sepsis. In addition, PEU significantly changed composition of the flora at both phylum and genus levels. Moreover, PEU significantly affected metabolism-related pathways such as tricarboxylic acid (TCA) cycle, fatty acid degradation, secondary bile acid biosynthesis, and others.

**Conclusions:** Taken together, PEU significantly inhibits LPS-induced inflammation *in vitro* and CLP-induced sepsis *in vivo*. Further, its anti-inflammatory effect is independent of the TLR4/myD88/NF-κB pathway. In addition, PEU improves the intestinal flora imbalance at the early stage of sepsis.

## Introduction

Sepsis is an acute systemic infectious disease with a rapid progress and a poor prognosis. Once not treated in time, it will develop into septic shock, resulting in organ failure and multiple organ dysfunction syndrome (MODS). A study has shown that 30% to 50% of populations in the United States suffer from sepsis yearly, and its incidence rate is increasing gradually [1]. In worldwide, more than five million patients die due to the related complications per year. The short-term mortality is up to about 20%∼25%[2]. Once septic shock occurs, the mortality will exceed 50% [3].

The essence of sepsis is an excessive inflammatory reaction triggered by PAMPs recognizing pattern recognition receptors (PRRs) such as TLR4, TLR9, and TLR2. Its original intention is to eliminate invading pathogens. However, under persistent stimulus of these PAMPs, proinflammatory cells are overactivated and large number of inflammatory mediators are secreted, thereby resulting in proinflammatory cytokine storms, which causes damages and dysfunctions of target tissues and organs [4].

As a crucial membranous PRR, TLR4 recognizes LPS, activating MyD88 and then translocating NF-κB into nucleus through a series of signal transduction pathways, which finally releases massive of inflammatory cytokines such as TNF-α and IL-6 to accelerate the development of sepsis. Thus, it may be a promising therapeutic strategy to search for reagents targeting TLRs to cure sepsis [5].

Besides sepsis-induced injuries in liver and kidney [6], intestinal barrier dysfunction is another conspicuous characteristic feature accompanied by an intestinal flora imbalance during sepsis [7, 8]. It has been confirmed that severity of sepsis is positively correlated to flora imbalance [9]. Although fecal microbiota transplantation (FMT) increases fecal flora probiotics and improves diversity of the intestinal flora structure in patients with mild sepsis [10], it is inclined to increase death risk due to occurrence of resistant pathogenic bacteria [11, 12].

Peucedanol (PEU), also called ulopterol, is a coumarin compound from the root of *Peucedanum japonicum*, which has been traditionally used to cure lung heat cough, wetness-heat-induced urodynia, sore and carbuncle [13]. Also, it has been found in *Melicope pteleifolia* (Champ. Ex Benth.) T. G. Hartley, which is commonly used for heat clearing and detoxification in the concept of traditional Chinese medicine (TCM) [14]. Its chemical name is 6-[(2R)-2, 3-dihydroxy-3-methylbutyl]-7-methoxychromen-2-one, and its molecular formula (CAS: 28095-18-3) is C_15_H_18_O_5_ (Molecular weight: 278.30). The chemical structure of PEU is shown in Figure 1.

**Figure 1.**
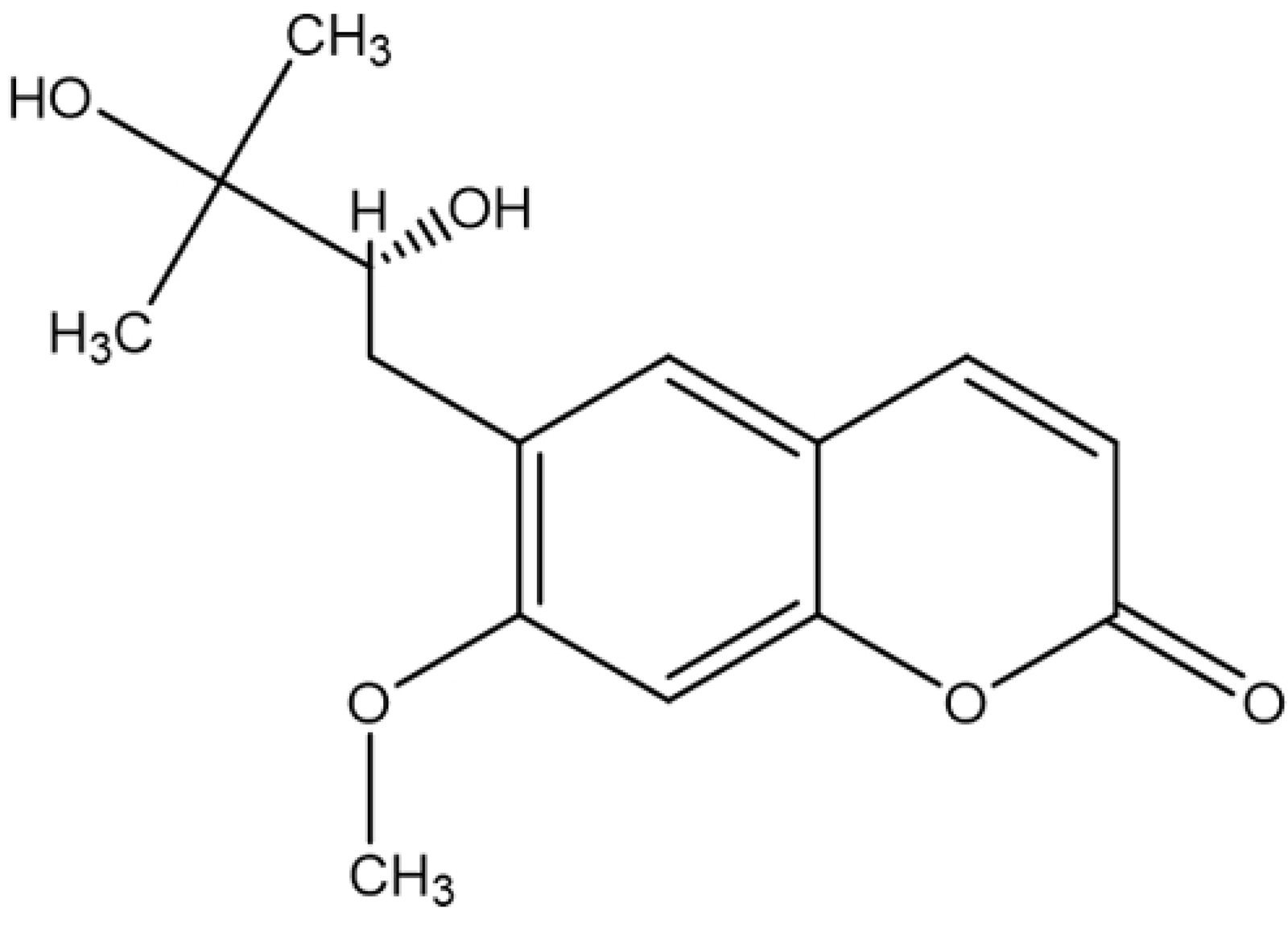
Chemical structure of PEU.

Nowadays, the pharmacological studies on PEU are few. An *in vitro* study showed that PEU was a non-competitive inhibitor of cytochrome3A4 (CYP3A4) and competitive inhibitors of CYP1A2 and CYP2D6, and its IC_50_ values for these three CYP450 enzymes were 6.03, 13.57, and 7.58 μM, respectively [15]. Further, PEU isolated from *Toddalia asiatica* (L.) Lam. significantly inhibited growth of some pathogenic bacteria including *Enterobacter aerogenes*, *Staphylococcus epidermidis*, *Shigella flexneri*, *Klebsiella pneumoniae* (ESBL-3967), *Escherichia coli* (ESBL-3984), and fungi [16].

Until now, few studies have documented the anti-inflammatory effect of PEU, let alone its mechanism. In the present study, ITC was used to measure the binding affinities of this compound to the PAMPs such as KLA, ODN 1826, and PGN. After that, the bioactivity of PEU was assessed in the related models *in vitro* and *in vivo*. Further, role of the TLR4/myD88/NF-κB pathway was investigated to explore its underlying mechanism in presence of CQ, a blocker of TLRs. Finally, its regulation for the intestinal flora during sepsis was analyzed using the 16S rRNA sequencing. This study will provide some cues for application of this compound in the treatment of sepsis-induced intestinal mucosal barrier dysfunction in the future.

## Materials and methods

### Experimental animal, cell line and main reagents

PEU (CAS: 20516-23-8, Batch No. BP1082, molecular weight: 264.27, formula: C_14_H_16_O_5_) (purity ≥ 98%) was purchased from Biochempartner Co., Ltd. (Shanghai, China). RAW264.7 cell line was provided by Procell Life Science & Technology Co., Ltd. (Wuhan, China). The male specific pathogen free (SPF) institute of cancer research (ICR) mice was obtained from Huabukang Biotechnology Co., Ltd. (Beijing, China). The product license number was SCXK (Beijing) 2019-0008, and the use license number was SYXK (Guizhou) 2021-0005. The animal experiments were conducted in accordance with the National Institutes of Health guide for the care and use of Laboratory animals (NIH Publications No. 8023, revised in 1978). The animal experiments were approved by the Experimental Animal Center of Guizhou University of TCM, and the ethics approval number was 20220095.

KLA (purity ≥ 90%) (CAS: 1246298-62-3, Batch No. T38635, molecular weight: 2306.84, formula: C_110_H_214_N_6_O_39_P_2_·xNH_3_) and PGN (CAS: 27814-48-8, Batch No. 78721-10MG-F, molecular weight: 119.0761, formula: C_3_H_5_NO_4_) from *Methanobacterium spp.* were purchased from Sigma-Aldrich Lab & Production Materials (City of Saint Louis, MO, USA). ODN 1826 (purity: 99.76%) (CAS: 202668-42-6, Batch No. HY-146245C, Sequence: d(P-thio) (T-C-C-A-T-G-A-C-G-T-T-C-C-T-G-A-C-G-T-T)) and chloroquine (CQ) (purity: 99.82%) (CAS: 54-05-7, Batch No. GC19549, molecular weight: 319.87, formula: C_18_H_26_ClN_3_) were provided by Glpbio (Montclair, CA, USA). The TLR4 primary antibody (Batch No.16c5074) was obtained from Affinity Biosciences (Changzhou, Jiangsu Province, China). The NF-κB primary antibody (Batch No. ab16502) was purchased from Abcam Inc. (Shanghai, China). The MyD88 primary antibody (Batch No. 10022637) was from Proteintech Group Inc. (Wuhan, China). The secondary anti-mouse antibody (Batch No. 23512) was obtained from Antibody & Life Science (Shenzhen, China). The secondary anti-rabbit antibody was purchased from Cell Signaling Technology (Shanghai, China). The mouse TNF-α (Batch No. 1217202) and IL-6 (Batch No. 1210602) ELISA kits were obtained from Dakewe Biotech Co., Ltd. (Shenzhen, China).

### ITC assay

KLA, ODN1826, and PGN were respectively dissolved in phosphate buffered saline (PBS), and then diluted with a PBS buffer containing 5% DMSO to reach 250 µM or 200 µM (pH7.4). After that, PEU was dissolved in DMSO, and then diluted with the PBS buffer supplemented with 5% DMSO (pH7.4) to reach a concentration of 10 µM. The pretreated solutions were filled in the injection wells (2 μl each well) at a room temperature. Equilibrated for 150 s, and then stirred at 750 rpm. Heat peaks were assayed by the Microcal PEAQ-ITC Control Software Update (version 1.30). The Microcal PEAQ-ITC Analysis Software (Malvern Panalytical, Malvern, UK) was used to generate thermodynamic parameters including equilibrium dissociation constant (Kd), enthalpy change (ΔH), and entropy change (ΔS).

### Cell viability assay

The RAW264.7 cells were cultured in Dulbecco’s modification of Eagle’s medium (DMEM) supplemented with 1% penicillin-streptomycin and 10% fetal bovine serum. After cultured in a 96-well plate (10^4^/well) for 12 h, and then replaced with serum-free medium. Respectively, 1 μl of PEU with various concentrations were added to co-culture for another 24 h in absence or presence of LPS (final concentration 200 ng/ml). After that, 10 μl of cell counting kit-8 (CCK-8) solution (Solarbio Life Sciences, Beijing, China) was added and co-cultured for 2 h. Finally, the optical density (OD) value of the solution was measured at 450 nm by using a Mutiskan Go Thermo microplate reader (Thermo Scientific, Massachusetts, USA).

### ELISA assay in vitro

The cells (10^5^/well) were seeded in a 6-well plate and cultured for 12 h. The cells were then treated in the absence or presence of CQ. In the absence of CQ, the cells were respectively treated with various concentrations of PEU (3.125-12.5 μg/ml) and DEX (4 μg/ml). In the presence of CQ, the cells were treated with PEU (12.5 μg/ml), CQ (2.5 μM), PEU plus CQ, and DEX (4 μg/ml). Subsequently, LPS was added to reach a final concentration of 200 ng/ml. Meanwhile, the vehicle was set parallelly. In the absence of CQ, 4, 8, 12 h after the LPS stimulation, 200 μl of the cell supernatants were collected to detect TNF-α and IL-6 levels using the ELISA method in accordance with the instructions of the manufacturer. Correspondingly, the cell precipitations were collected at 8 h for western blotting assay. In the presence of CQ, 8 h after the LPS stimulation, the cell supernatants and precipitations were respectively collected for the ELISA and western blotting.

### Western blotting assay

The collected cell precipitations were lysed by boiling 1 × loading buffer (Beyotime Biotechnology, Haimen, Jiangsu, China) and then ultrasonicated on ice for 30 seconds by using a LANYI-650Y ultrasonic cell disrupter system (Lanyi Instrument Co., Ltd., Shanghai, China) at 22 ± 1 KHz and 80 w. Centrifuged at 13,000 ⊆ *g* for 5 min at 4°C to harvest the supernatants. Next, the total proteins were separated by 10% sodium dodecyl sulfate polyacrylamide gel electrophoresis (SDS-PAGE) followed by semi-dry transfer onto 0.22-μm polyvinylidene difluoride (PVDF) membranes. Blocked with 5% fat-free milk dissolved in tris buffered saline with Tween20 (TBST). After that, added primary anti-TLR4 (1: 1000), -MyD88 (1: 2000), -NF-κB (1: 2000) antibodies respectively to co-culture for 2.0 h at room temperature. Added secondary anti-mouse antibody (1: 6000) or anti-rabbit antibody (1: 6000) respectively to co-culture for another 1 h. Imaged by enhanced chemiluminescence (ECL) with a ChemiDoc^TM^ Imaging System (BIO-RAD, Hercules, CA, USA). β-actin was selected as an internal control.

### Establishment of CLP-induced sepsis mouse model and treatments

The male ICR mice weighing 20-22 g were randomly divided into a sham group, a model group, a dexamethasone (DEX) group (4 mg/kg, i.g), and three PEU groups (6.25, 12.5, 25 mg/kg, i.g, 12 animals in each group) (PEU dissolved in 5% CMC-Na). The mice received various treatments once a day and it lasted five days continuously. Four hours after the last treatments, the animals received the CLP surgeries except for the sham ones. Six hours post the surgeries, the peripheral blood samples were collected by sequential bleeds and then centrifuged to harvest the sera for the ELISA assay (n = 8). Recorded the death of all the animals every day within 7 days. At the end of the experiment, the survival curves of different groups were plotted to assess the anti-infective effect of PEU. The small intestine samples were collected for the pathological examination after the animals were sacrificed.

In the presence of CQ, the male ICR mice were randomly separated into a sham group, a model group, a PEU group (25 mg/kg, i.g), a CQ group (20 mg/kg, i.g) (dissolved in 5% CMC-Na), and a PEU plus CQ group (n = 12). The animals received treatments once a day and it lasted five days continuously. Four hours after the last treatments, the CLP surgeries were performed except for the sham ones. Six hours after the surgeries, the peripheral blood samples were collected using terminal bleeds to harvest the sera to determine the serum TNF-α and IL-6 levels (n = 6). At the end of the experiment (within 168 h), the survival curves were plotted as described before.

### Pathological examination of small intestine

The collected small intestine samples were fixed in 4% paraformaldehyde for 24 h. They were dehydrated and embedded in paraffins. Next, the paraffin sections were cut into 2 μm-thick sections consecutively followed by staining with hematoxylin and eosin (H&E). The pathological changes especially edema, inflammation, and hemorrhage in the tissues were observed under a DM750 microscope and imaging system (Leica, Weztlar, Germany).

### Immunofluorescence staining assay

The small intestine paraffin sections were deparaffinized and subjected to antigen repairing using sodium citrate antigen repair solution in a microwave oven. Cooled to room temperature and washed three times with PBS. Dried the slices and added 5% BSA to block for 30 min and then incubated with anti-rabbit Bmi1 (1: 50) (Batch No. PB0102, Boster Biological Technology, Wuhan, China) or anti-rabbit Lgr5 (1: 50) (Batch No. A00239-2, Boster Biological Technology, Wuhan, China) antibody overnight at 4℃ in a wet box. Washed with PBS three times. Added fluorescence-labelled secondary antibody and incubated in the dark at 37°C for 50 min. Subsequently, washed three times, and then added anti-fluorescence quencher containing DAPI (Thermo Fisher Scientific, Waltham, MA, United States) for sealing. Imaged and observed the images under an inverted fluorescence microscope.

### Feces collection and 16S rRNA sequencing

A total of 50 male SPF ICR mice were randomly separated into a blank group, an LPS-treated group, two PEU-treated groups (12.5 and 25 mg/kg), and a dexamethasone (DEX) group (8 mg/kg, i.g) (n = 10). The mice in the groups received various treatments for continuous five days and once a day as before. Four hours after the last administrations, LPS solutions dissolved in normal saline (NS) were injected intraperitoneally into the mice (20 mg/kg, 0.1 ml/mouse) except for the blank ones. Correspondingly, the mice in the blank group were injected with the equal volumes of NS.

The feces samples were collected in sterile saline in 1.5-ml sterile tubes 6 h after the LPS injections. The 16S rRNA sequencing was performed as follows. Total genome DNAs were extracted from the feces samples by using cetyltrimethylammonium bromide (CTAB). The concentration and purity of the extracted DNAs were determined by 1% agarose gels electrophoresis. After that, 16S rRNA genes of distinct regions of 16S V3-V4 were amplified by using a forward primer (338F) “5’-ACTCCTACGGGAGGCAGCAG-3’” and a reverse primer (806R) “5’-GGACTACHVGGGTWTCTAAT-3’. The PCR reaction system included 10 ng template DNA, and 15 μl Phusion®High-Fidelity PCR Master Mix (New England Biolabs, Beijing, China), and 0.2 μM forward and reverse primers. The thermal cycling parameters included 30 cycles of denaturation at 98°C for 30 s, annealing at 50°C for 30 s, and extension at 72°C for 30 s. The PCR products were then mixed with IX loading buffer containing SYB green (v/v, 1:1) and examined by 2% agarose gel. Next, the mixture was purified by using a Qiagen Gel Extraction Kit (Qiagen, Dusseldorf, Germany).

A TruSeq® DNA PCR-Free Sample Preparation Kit (Illumina, San Diego, CA, USA) was used to generate the sequencing libraries according to the manufacturer’s recommendations. The quality of the library was assessed by the Qubit@2.0 Fluorometer (Thermo Scientific) and Agilent Bioanalyzer 2100 system. The library was sequenced to generate 250-bp paired-end reads by an Illumina NovaSeq platform. The sequences possessing quality score less than 20, fragment length shorter than 200 bp, or ambiguous bases and mismatch primer sequences were removed. The qualified reads were separated and trimed by the Illumina Analysis Pipeline (Version 2.6) based on the sample-specific barcode sequences. The vsearch software (Version 2.8.1) was then used to analyze the dataset. At a similarity level of 97%, the sequences were clustered into operational taxonomic units (OTUs) to generate rarefaction curves and calculate the richness and diversity indices. The similarity between different samples were examined by the clustering analyses and principal component analysis (PCA) based on the information of each sample.

### Data presentation and statics

The data were expressed as mean ± standard deviation. One-way analysis of variance (ANOVA) was used to compare statistical significances among groups. A *p* value of less than 0.05 is regarded as significant.

## Results

### PEU has moderate binding to KLA and ODN 1826, but not PGN

The dissociation constant (Kd) values between PEU and KLA and ODN 1826 were respectively 177e^-6^ ± 2.36e^-3^ mol/1 and 18.9e^-6^ ± 29.8e^-6^ mol/l, while it had no significant binding to PGN (Figure 2). Further, PEU had a stronger affinity to ODN1826 compared to KLA (Figure 2).

**Figure 2.**
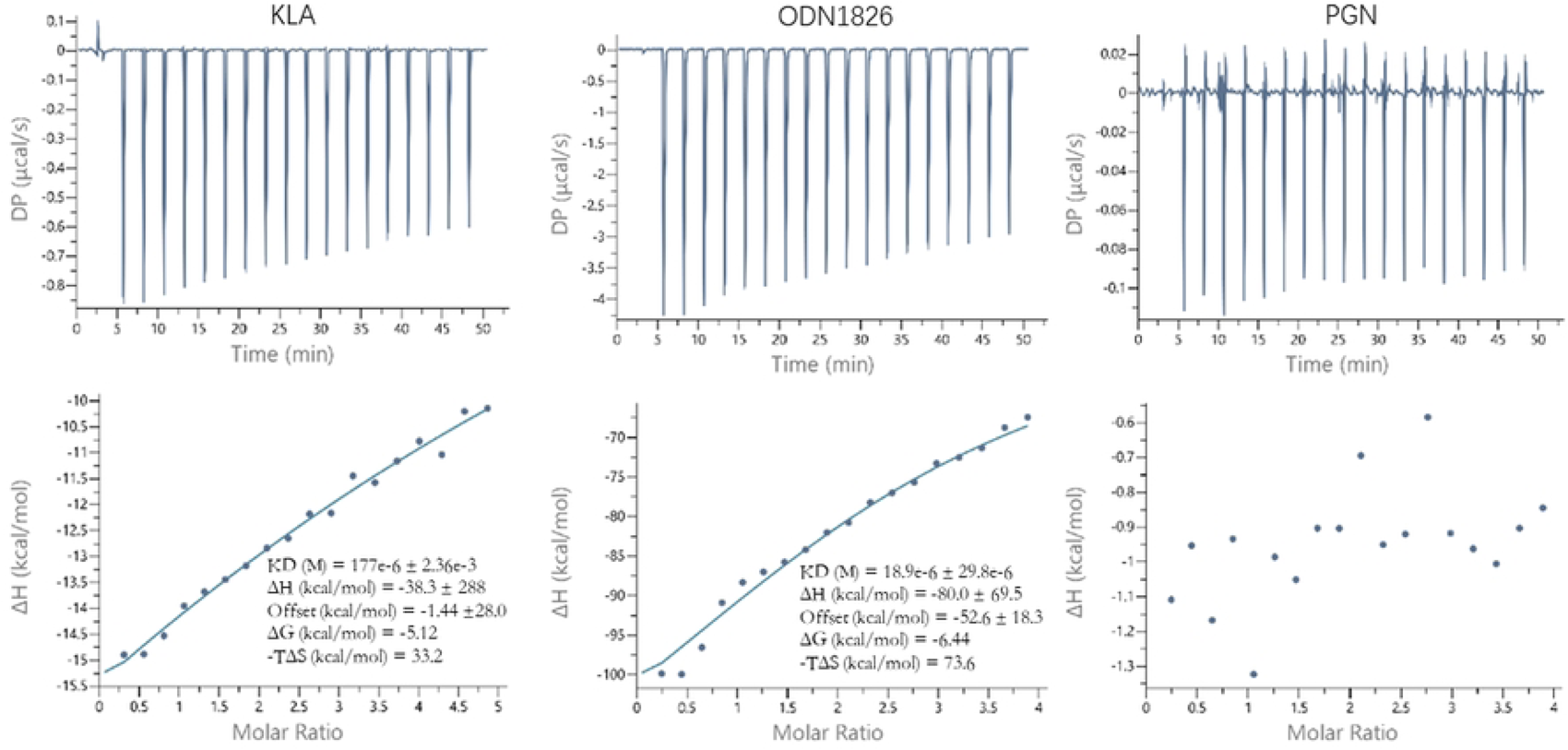
ITC analysis of binding affinities of PEU to KLA, ODN1826, and PGN. KLA, ODN1826, and PGN were respectively dissolved to reach 250 µM or 200 µM. Meanwhile, PEU was dissolved with DMSO, and then diluted with PBS buffer to reach 10 µM. 2 μl of pretreated protein solutions were filled in the injection cells and equilibrated. After that, the Microcal PEAQ-ITC analysis software was used to assay the binding affinities of PEU to the three proteins.

### PEU inhibits LPS-induced inflammation in vitro

The CCK-8 showed that PEU > 25 μg/ml significantly inhibited the cell growth, and PEU ≤ 12.5 μg/ml had no significant toxicity for the RAW264.7 cells (Figure 3A). Further, PEU at 3.125-12.5 μg/ml significantly promoted the LPS-treated cell growth compared to the LPS treatment alone (*p* < 0.05). The cell proliferations were 96.4%, 97.3%, 97.8% for 3.125, 6.25, and 12.5 μg/ml PEU, respectively (Figure 3B).

**Figure 3.**
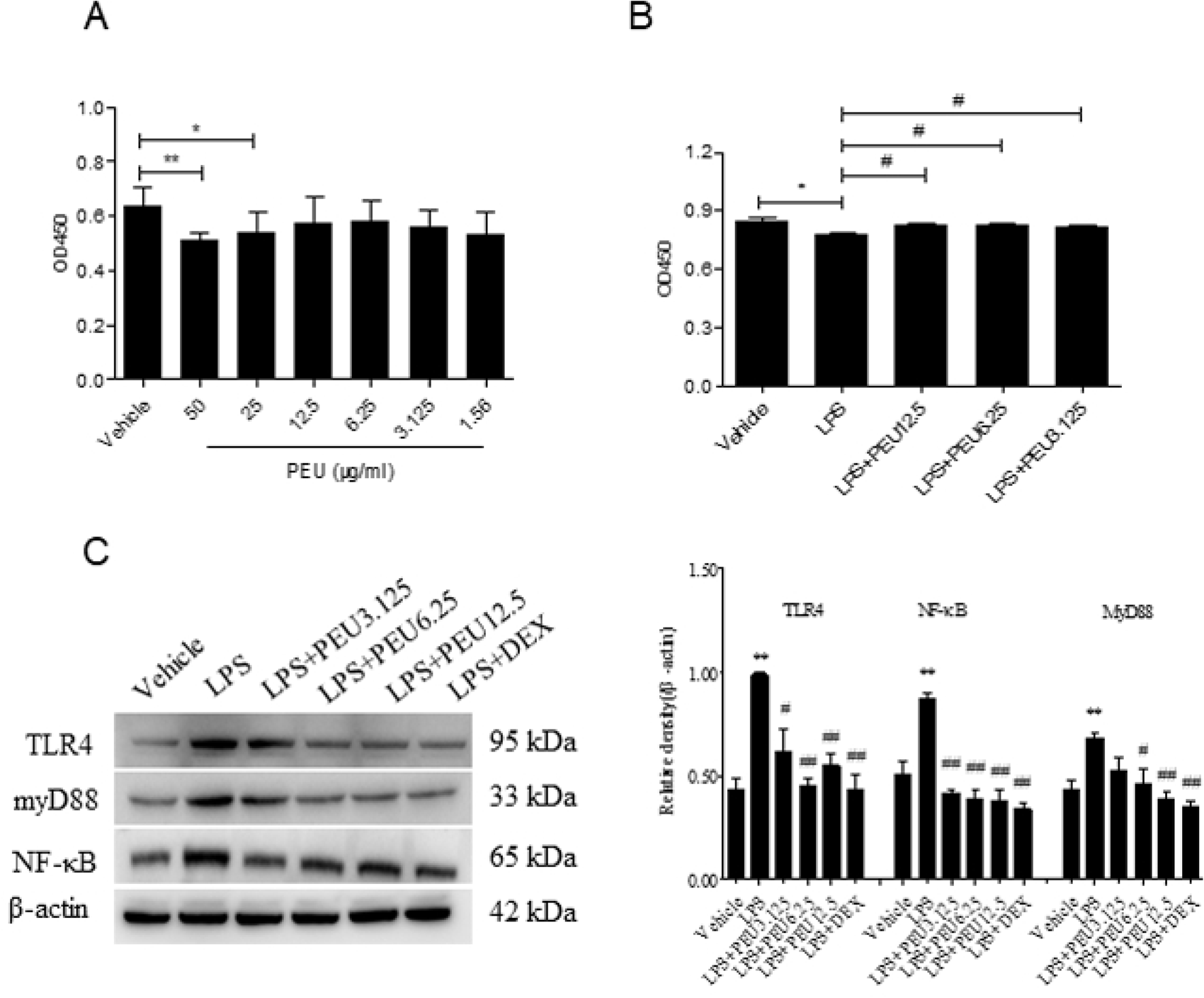
PEU inhibits LPS-induced inflammation *in vitro*. **A and B**, Cell toxicity of PEU in RAW264.7 cells in absence or presence of LPS (x̅ ±s, n=3). \**p*<0.05, \*\**p*<0.01 vs. Vehicle. The cells were respectively co-cultured with 1 μl of various concentrations of PEU (1.56, 3.125, 6.25, 12.5, 25, 50 μg/ml) for 24 h followed by adding of 10 μl CCK-8 for another 2 h in serum-free medium. The OD value of the solution was measured at 450 nm by using a microplate reader; **C,** PEU downregulates the protein expressions of TLR4, myD88, and NF-κB in LPS-stimulated cells (x̅ ±s, n=3). \**p*<0.05, \*\**p*<0.01 vs. Vehicle; ^#^*p*<0.05, ^##^*p*<0.01 vs. LPS. The cells were treated with PEU (3.125-12.5 μg/ml) in presence of 200 ng/ml of LPS for 4, 8, and 12 h, respectively. Collected the cell precipitations and lysed. The total proteins were then separated by using 10% SDS-PAGE. Next, western blotting was performed to measure the expressions of TLR4, myD88, and NF-κB proteins in the cell precipitations. Finally, imaged by using ECL in an Imaging System.

The ELISA result demonstrated that the LPS treatment significantly increased the supernatant TNF-α and IL-6 levels compared to the vehicle at 4, 8, 12 h respectively (*p* < 0.05) (Table 1 and 2). Compared with the LPS treatment, PEU at 6.25 and 12.5 μg/ml remarkably downregulated the increased supernatant TNF-α and IL-6 levels (Table 1 and 2).

**Table 1.** Supernatant TNF-α level (pg/ml) at regular time points (x̅ ±s, n = 3)

| Treatment | Dose<br>($\mu\text{g/ml}$) | Time (Hours) | | |
| --- | --- | --- | --- | --- |
|  |  | 4 | 8 | 12 |
| Vehicle | — | 25.5±2.8 | 32.5±4.2 | 45.2±9.4 |
| LPS | — | 1339.8±120.0* | 1698.9±132.0* | 2601.2±501.4* |
| LPS+DEX | 5 | 554.7±98.5*<sup>##</sup> | 735.4±133.1*<sup>##</sup> | 965.2±238.8*<sup>##</sup> |
| LPS+Low-dose PEU | 3.125 | 1375.0±179.0* | 1932.3±323.1* | 2659.5±700.9* |
| LPS+Middle-dose PEU | 6.25 | 801.9±130.3*<sup>##</sup> | 1113.2±200.2*<sup>#</sup> | 1604.5±337.6*<sup>#</sup> |
| LPS+High-dose PEU | 12.5 | 745.5±137.0*<sup>##</sup> | 1047.4±254.9*<sup>#</sup> | 1446.5±345.1*<sup>#</sup> |

**Table 2.** Supernatant IL-6 level (pg/ml) at regular time points (x̅ ±s, n = 3)

| Treatment | Dose<br>($\mu\text{g/ml}$) | Time (Hours) | | |
| --- | --- | --- | --- | --- |
|  |  | 4 | 8 | 12 |
| Vehicle | — | 29.3±2.4 | 35.8±3.3 | 44.6±6.8 |
| LPS | — | 1428.1±183.5* | 1700.4±145.7* | 1989.9±380.7* |
| LPS+DEX | 5 | 430.3±53.0*<sup>##</sup> | 610.0±97.3*<sup>##</sup> | 948.8±100.1*<sup>#</sup> |
| LPS+Low-dose PEU | 3.125 | 1601.0±300.2* | 1836.8±450.1* | 2166.0±517.2* |
| LPS+Middle-dose PEU | 6.25 | 727.9±126.5*<sup>##</sup> | 953.6±230.5*<sup>##</sup> | 1408.6±298.2* |
| LPS+High-dose PEU | 12.5 | 554.4±97.7*<sup>##</sup> | 732.9±99.1*<sup>##</sup> | 1103.8±128.5*<sup>#</sup> |

\* $p < 0.01$ vs. Vehicle; # $p < 0.05$ , <sup>##</sup> $p < 0.01$ vs. LPS. The cells were respectively co-cultured with 1 $\mu\text{l}$ of PEU (3.125, 6.25, 12.5 $\mu\text{g/ml}$ ) in presence of LPS with a final concentration of 200 ng/ml. Co-cultured and collected the supernatants for 4, 8, and 12 h, respectively. Finally, detected the OD450 by using a microplate reader.

The western blotting assay showed that the protein expressions of TLR4, myD88, and NF-κB in the cells were significantly elevated 8 h after the LPS treatment compared to the vehicle (*p* < 0.05) (Figure 3C). It revealed that PEU markedly downregulated the increased expressions of these three proteins in the LPS-treated cells (Figure 3C), suggesting a significant inhibitory effect of this compound on the TLR4/myD88/NF-κB pathway.

### PEU protects CLP-induced sepsis mice

After received the CLP surgeries, the animals began to die and the death time of major animals was from 24 to 48 h. All the animals in the CLP group died within 72 h, suggesting the establishment of CLP-induced sepsis mouse model (Figure 4A). It showed that the PEU pretreatments (12.5 and 25 mg/kg) for five days significantly increased the survival rate of the sepsis mice compared to the model (*p* < 0.05). The survival rates in the PEU groups were respectively 75% for 25 mg/kg and 66.7% for 12.5 mg/kg, suggesting a significant protective effect of this compound for the CLP-induced sepsis mice (Figure 4A).

**Figure 4.**
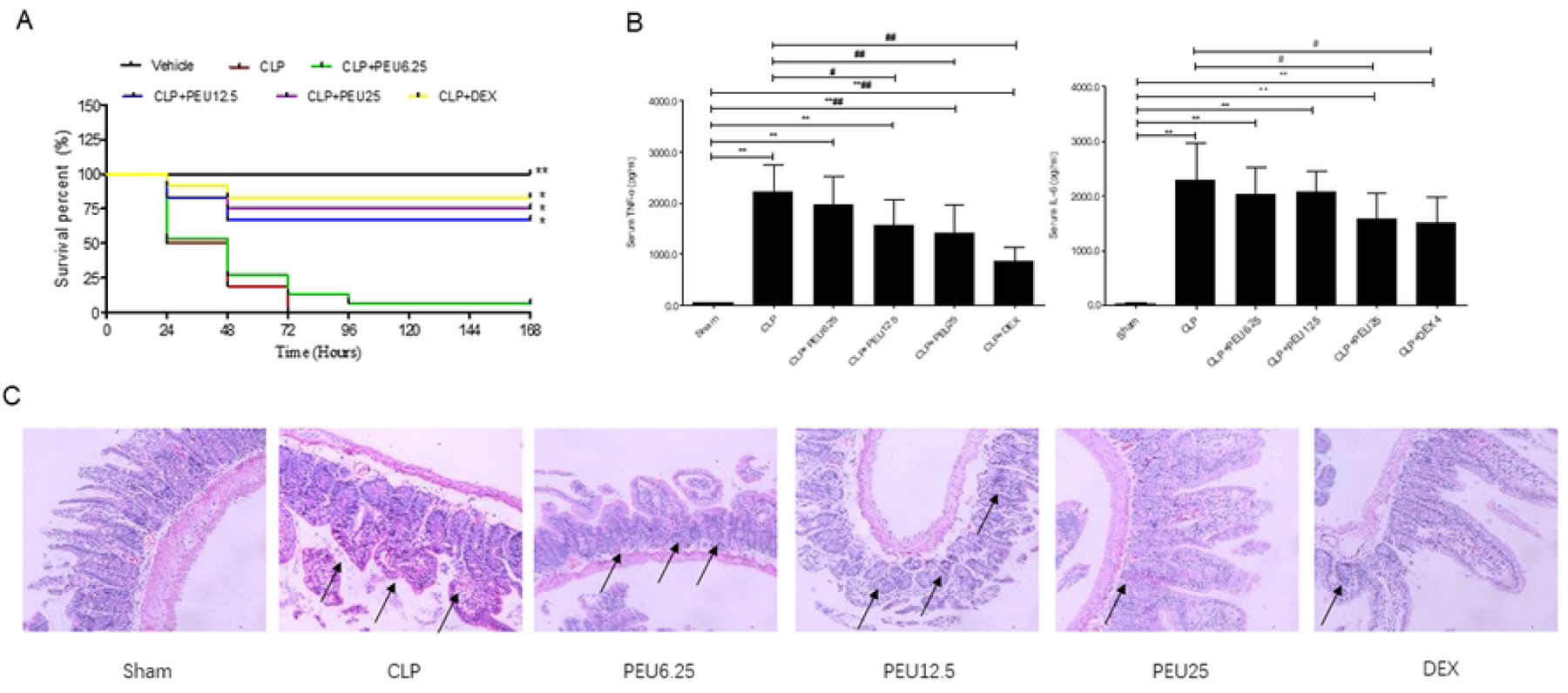
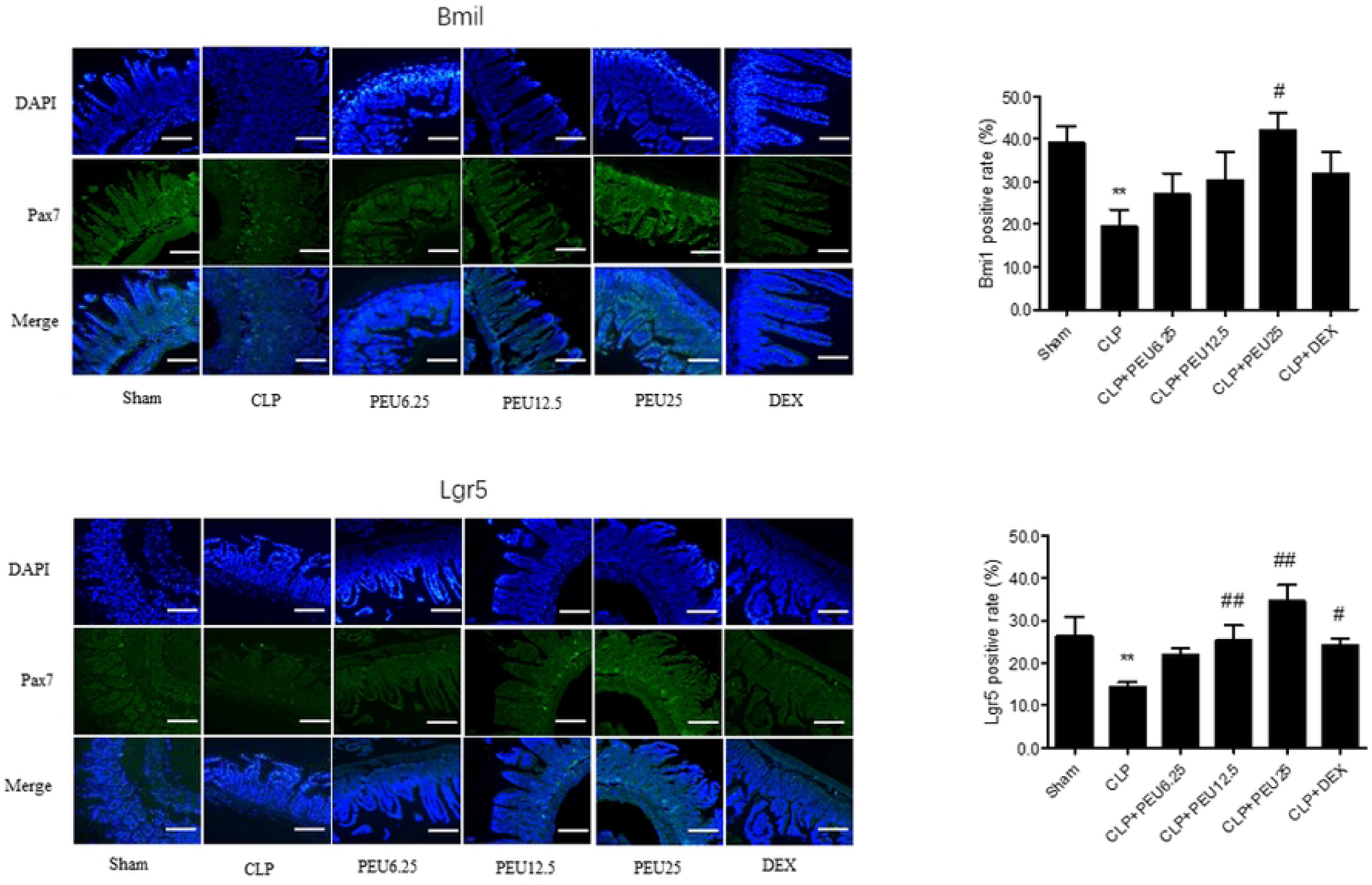
PEU protects CLP-induced sepsis mice. **A**, PEU increases the survival rate of the sepsis mice. \**P*<0.05, \*\**P*<0.01 vs. CLP. Before the CLP surgeries, the mice received various treatments for five days and once a day. 4 h post the last treatments, the CLP surgeries were performed in all the animals except for the sham ones. After the CLP surgeries, deaths of the animals in groups were recorded within 168 h and plotted to draw survival curves to evaluate the anti-infection of the reagents. The differences among groups were examined by Kaplan-Meier method; **B,** PEU decreases serum TNF-α and IL-6 levels in sepsis mice (x̅ ±s, n = 8). \*\**P*<0.01 vs. Sham ^#^*P*<0.05, ^##^*P*<0.01 vs. CLP. Six hours post the surgeries, the peripheral blood samples were collected by sequential bleeds to determine serum TNF-α and IL-6 levels by ELISA; **C,** PEU treatment attenuates pathological injuries of small intestines of sepsis mice (H&E). The collected small intestines were fixed, dehydrated and embedded. The tissue samples were then cut into 2 μm-thick sections and stained with H&E. The pathological changes especially edema, inflammation, and hemorrhage in the tissues were observed under an optical microscope; **D**, PEU upregulates Bmi1 and Lgr5 in the small intestines (x̅ ±s, n = 3). \*\**P*<0.01 vs. Sham ^#^*P*<0.05, ^##^*P*<0.01 vs. CLP. The small intestine paraffin sections were deparaffinized and subjected to antigen repairing. Then the Bmi1 or Lgr5 antibody was co-cultured overnight at 4℃ in a wet box. After that, fluorescence-labelled secondary antibody Added and incubated in the dark at 37°C for 50 min. Then added anti-fluorescence quencher containing DAPI, imaged and observed the images under an inverted fluorescence microscope.

Compared with the sham, the CLP surgery remarkably increased the serum TNF-α and IL-6 levels (*p* < 0.05) (Figure 4B). The PEU pretreatment markedly reduced the increased serum TNF-α and IL-6 levels compared to the CLP treatment alone (*p* < 0.05) (Figure 4B), which was some consistent with the ELISA results *in vitro*.

It was noted that the CLP surgery resulted in the disordered arrangements, destroyed structures, obvious swelling, even disintegration and death of some intestinal goblet cells companied by significant inflammatory infiltrations (Figure 4C). The PEU pretreatments (12.5-25 mg/kg) in varying degrees improved damaged intestinal structure, including reducing cellular degeneration and necrosis, alleviating inflammatory infiltration and congestion (Figure 4C).

Further, the CLP significantly downregulated the positive expressions of Bmi1 and Lgr5 proteins in the intestines compared to the sham (*p* < 0.05). Compared with the CLP, the PEU pretreatments (12.5 and 25 mg/kg) significantly upregulated the decreased positive expressions of Bmi1 and Lgr5 proteins in the intestine tissues (Figure 4D).

### PEU exerts anti-inflammatory effects independent of TLR4/myD88/NF-κB pathway

*In vitro*, the LPS treatment significantly increased the supernatant TNF-α and IL-6 levels compared to the vehicle 8 h after the LPS stimulation (*p* < 0.01). Compared with the LPS treatment, the PEU or CQ alone significantly lowered the supernatant TNF-α and IL-6 (*p* < 0.01). Further, the PEU plus CQ further downregulated the supernatant TNF-α and IL-6 significantly compared to the PEU or CQ alone (*p* < 0.05) (Figure 5A). In addition, the inhibitory effect of PEU on the protein expressions of TLR4, myD88, and NF-κB were not significantly abolished 8 h after the LPS treatment in the presence of CQ (Figure 5B). It suggested PEU attenuated LPS-induced cellular inflammatory responses independent of the TLR4/myD88/NF-κB pathway. In other words, other signal pathways might participate in the *in vitro* anti-inflammatory event of PEU.

**Figure 5.**
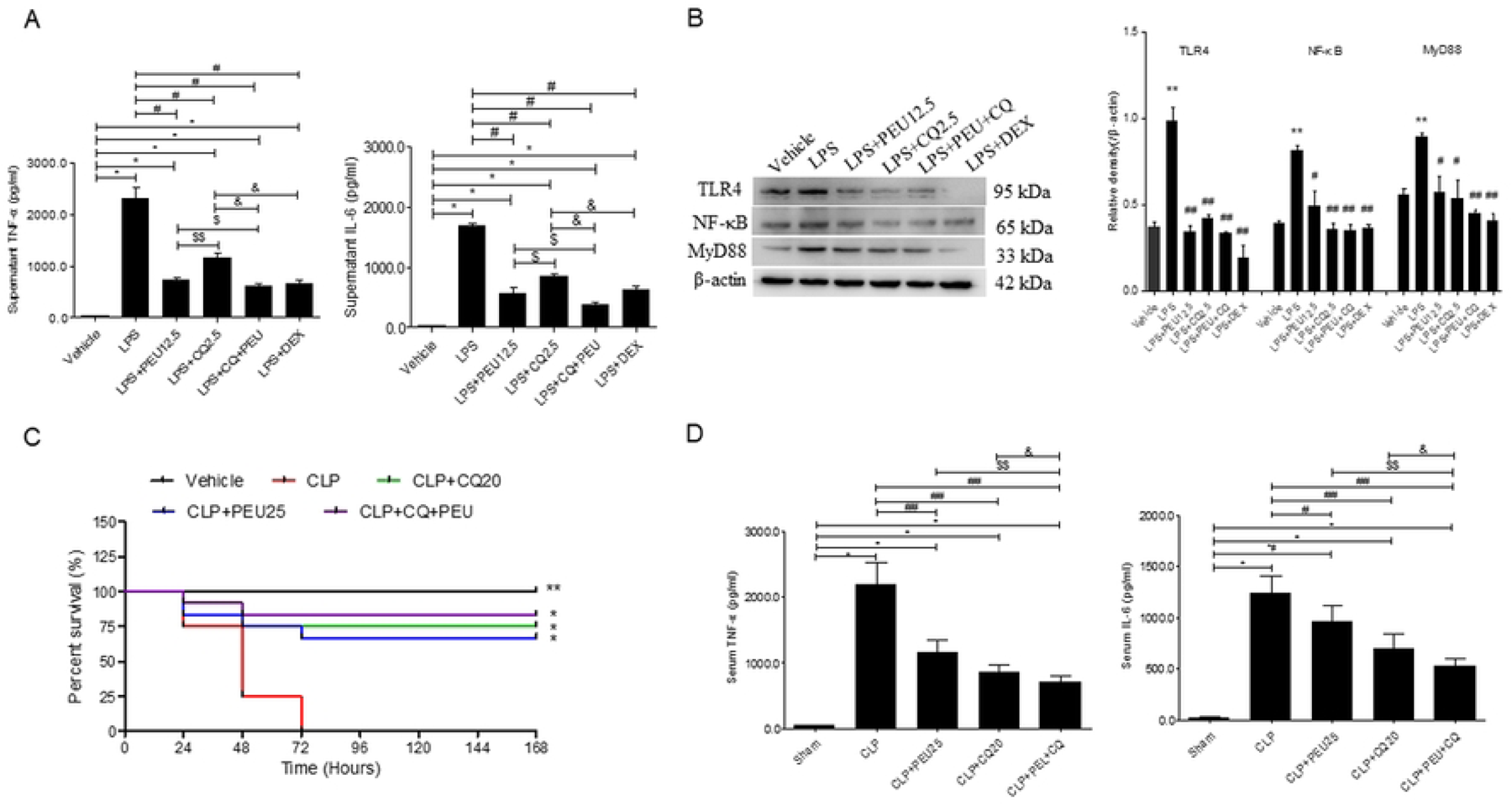
Role of TLR4/myD88/NF-κB pathway in anti-inflammatory effect of PEU. **A**, Supernatant TNF-α and IL-6 levels in presence of CQ (x̅ ±s, n=3). \**p*<0.01 vs. Vehicle; ^#^*p*<0.01 vs. LPS; ^$^*p*<0.05, ^$$^*p*<0.01 vs. LPS+PEU; ^&^*p*<0.01 vs. LPS+CQ. The LPS-stimulated cells were respectively treated with CQ (2.5 μM), PEU (12.5 μg/ml), CQ plus PEU, and DEX (5 μg/ml) for 8 h. The cell supernatants and precipitations were collected for the ELISA and western blotting assays. The supernatant TNF-α and IL-6 levels were determined using ELISA; **B**, The expressions of TLR4, myD88, and NF-κB in LPS-treated cells in presence of CQ (x̅ ±s, n=3). \**p*<0.05, \*\**p*<0.01 vs. Vehicle; ^#^*p*<0.05, ^##^*p*<0.01 vs. LPS. The collected precipitations were lysed for the western blotting assay; **C**, Survival of sepsis mice in presence of CQ. \**p*<0.05, \*\**p*<0.01 vs. CLP (Kaplan-Meier). The mice were pretreated with PEU, CQ, PEU plus CQ for five days and once a day. Four hours post the last treatments, the animals were received the CLP surgeries. The death of the animals within 168 h were recorded to plot survival curves; **D,** Serum TNF-α and IL-6 levels in sepsis mice in presence of CQ (x̅ ±s, n = 6). \*\**p*<0.01 vs. sham ^#^*p*<0.05, ^##^*p*<0.01 vs. CLP. The CLP-induced sepsis animals were pretreated with PEU, CQ, PEU combined with CQ as before. 6 h after the CLP surgeries, the peripheral blood samples were collected to detect the serum TNF-α and IL-6 levels by ELISA.

*In vivo*, the PEU or CQ alone significantly protected the sepsis mice, with the survival rates of 75% and 75%, respectively (*p* < 0.05). However, the CQ plus PEU didn’t significantly increase the survival rate compared to the CQ or PEU alone (*p* > 0.05) (Figure 5C). Compared with the CLP, the PEU or CQ alone significantly reduced the serum TNF-α and IL-6 levels of the sepsis mice (*p* < 0.05). Moreover, PEU plus CQ further significantly downregulated the two cytokines compared to the PEU or CQ alone (*p* < 0.05), which was also consistent with the ELISA result *in vitro* (Figure 5D). It suggested that the PEU pretreatment protected the CLP-induced sepsis mice by decreasing the serum TNF-α and IL-6 levels. Also, PEU reduced the CLP-induced inflammatory responses *in vivo* independent of the TLR4/myD88/NF-κB pathway as well as *in vitro*.

### Effects of PEU on α and β diversities of intestinal flora

No significant differences were present in α diversity indices such as Chao1, Shannon, and Simpson of the intestinal flora between the blank group and the LPS group (*p* > 0.05). All these three indices were lower in the DEX-treated group (8 mg/kg) than those in the LPS-treated group (*p* < 0.05). Compared with the LPS treatment, PEU (12.5 mg/kg) markedly increased the Chao1 index (*p* < 0.05), whereas it had no significant influences on the Shannon and Simpson indices (*p* > 0.05) (Figure 6A). It suggested that PEU significantly increased species richness of the intestinal flora during sepsis.

**Figure 6.**
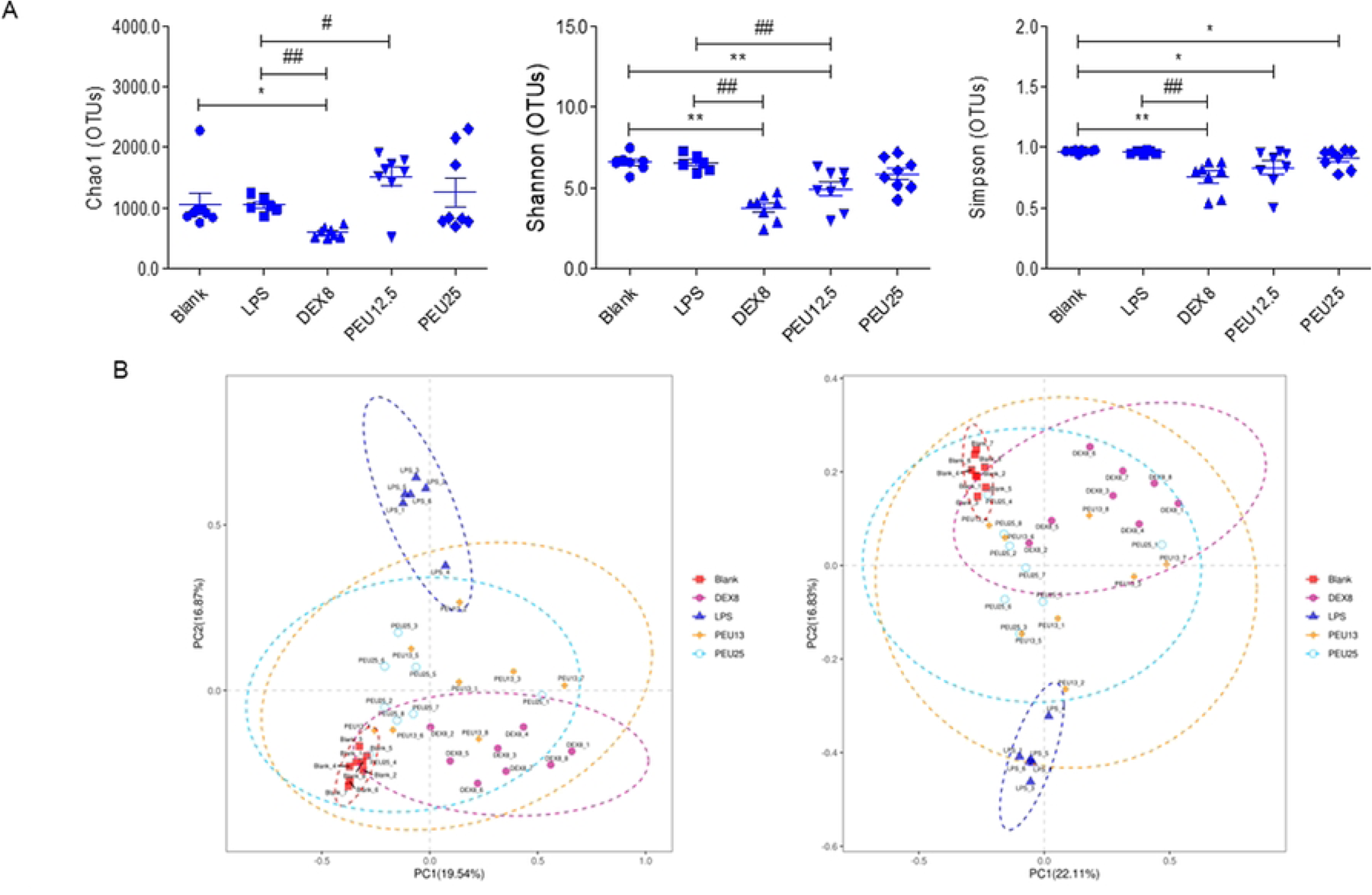
α and β diversities of the intestinal flora of the sepsis mice. **A**, α diversity of intestinal flora after treatments (x̅ ±s, n=5-8). \**p*<0.05, \*\**p*<0.01 vs. Blank; ^#^*p*<0.05, ^##^*p*<0.01 vs. LPS. The mice were respectively treated with low-dose PEU (12.5 mg/kg), high-dose PEU (25 mg/kg), and DEX (8 mg/kg) for five days and once a day. Four hours after the treatments, LPS were intraperitoneally injected into the mice (20 mg/kg) except for the vehicle controls. The feces samples were collected six hours after the LPS injections. The 16S rRNA sequencing was then performed. The α diversity indices including Chao-1, Shannon, and Simpon were detected to compare the differences among the groups; **B**, β diversity of intestinal flora after treatments. The PCoA assay was used to clarify sample separation and a clear clustering in the groups, and tree species were used to assay similarity of intragroup bacterial populations in each group.

The principal coordinates analysis (PcoA) and similarity analysis revealed good sample separation and a clear clustering in the groups for β diversity of the intestinal flora (Figure 6B).

### Changes in flora composition and metabolism pathways during sepsis

A total of 28 bacteria were significantly changed after the treatments at the phylum level (*p* < 0.05). For example, the proportions of *Bacteroidota*, *Deferribacterota*, and *Campilobacterota* were higher, and the proportions of *Firmicutes*, *Desulfobacterota*, and *Actinobacteriota* were lower in the PEU-treated groups than those in the LPS-treated group, respectively (Figure 7A).

**Figure 7.**
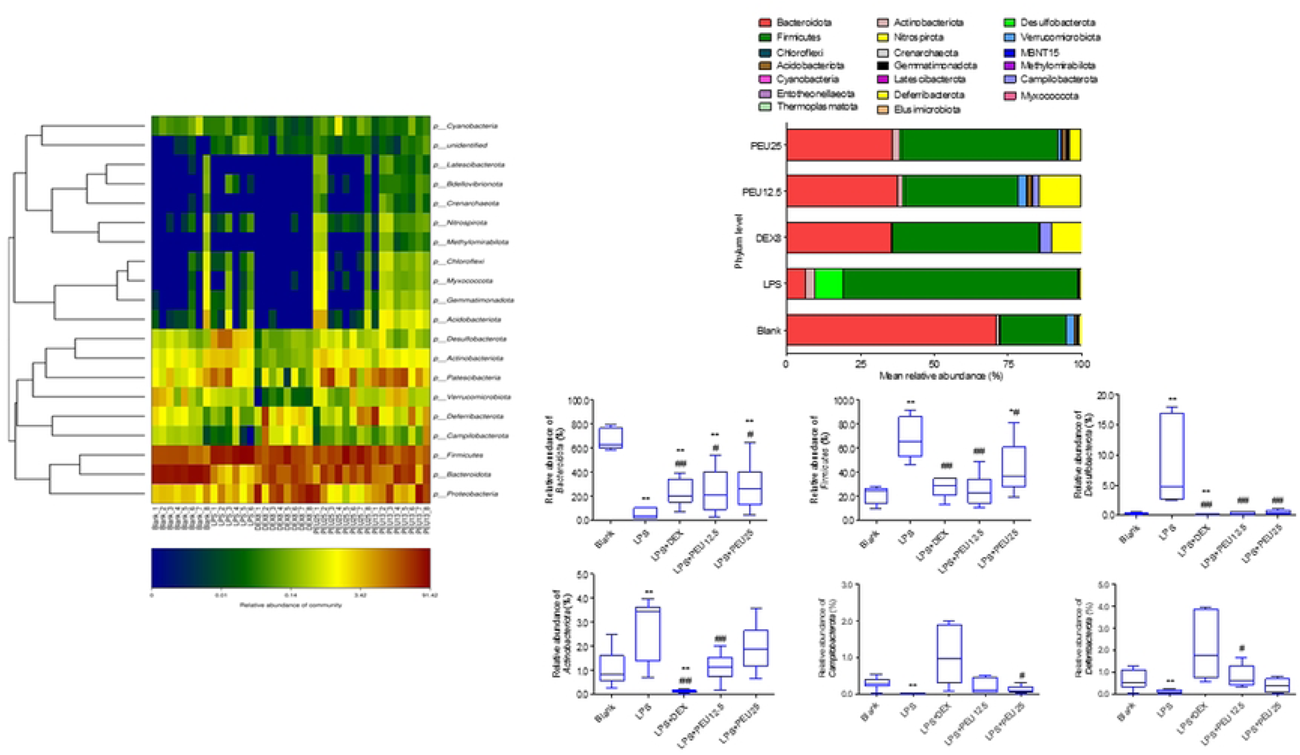
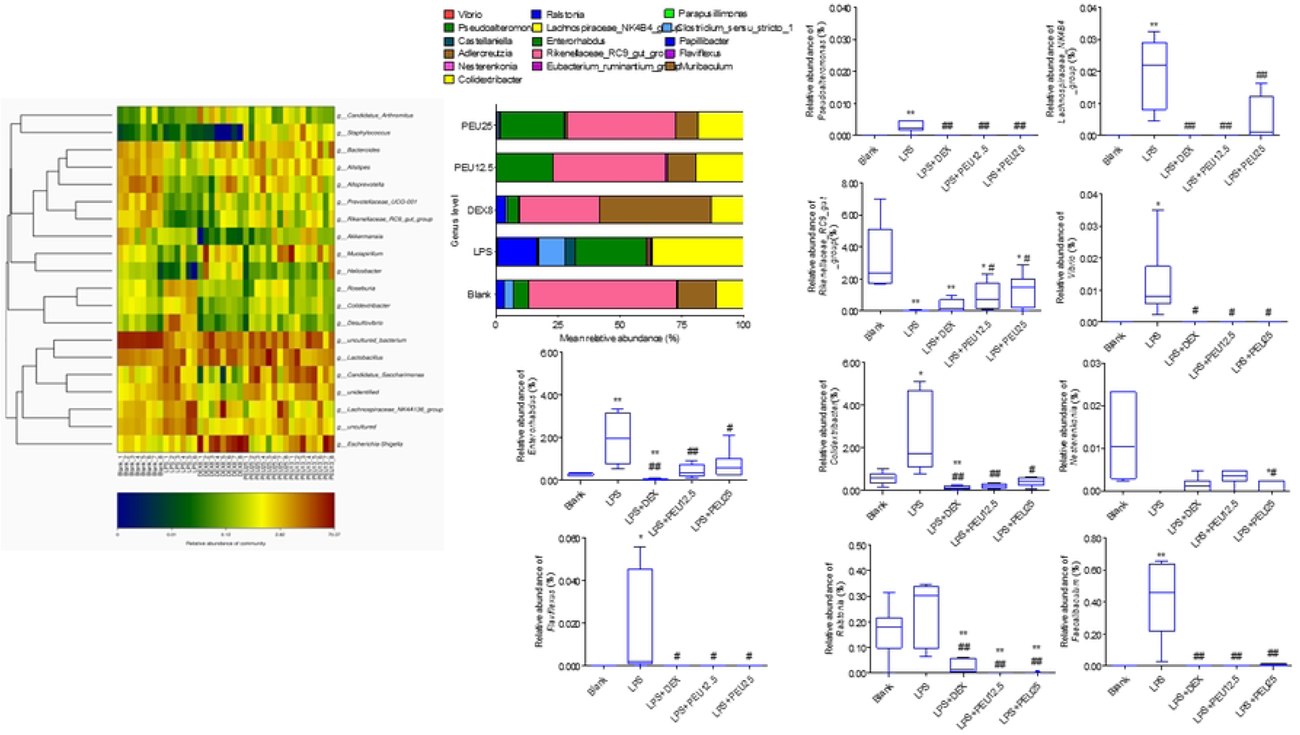
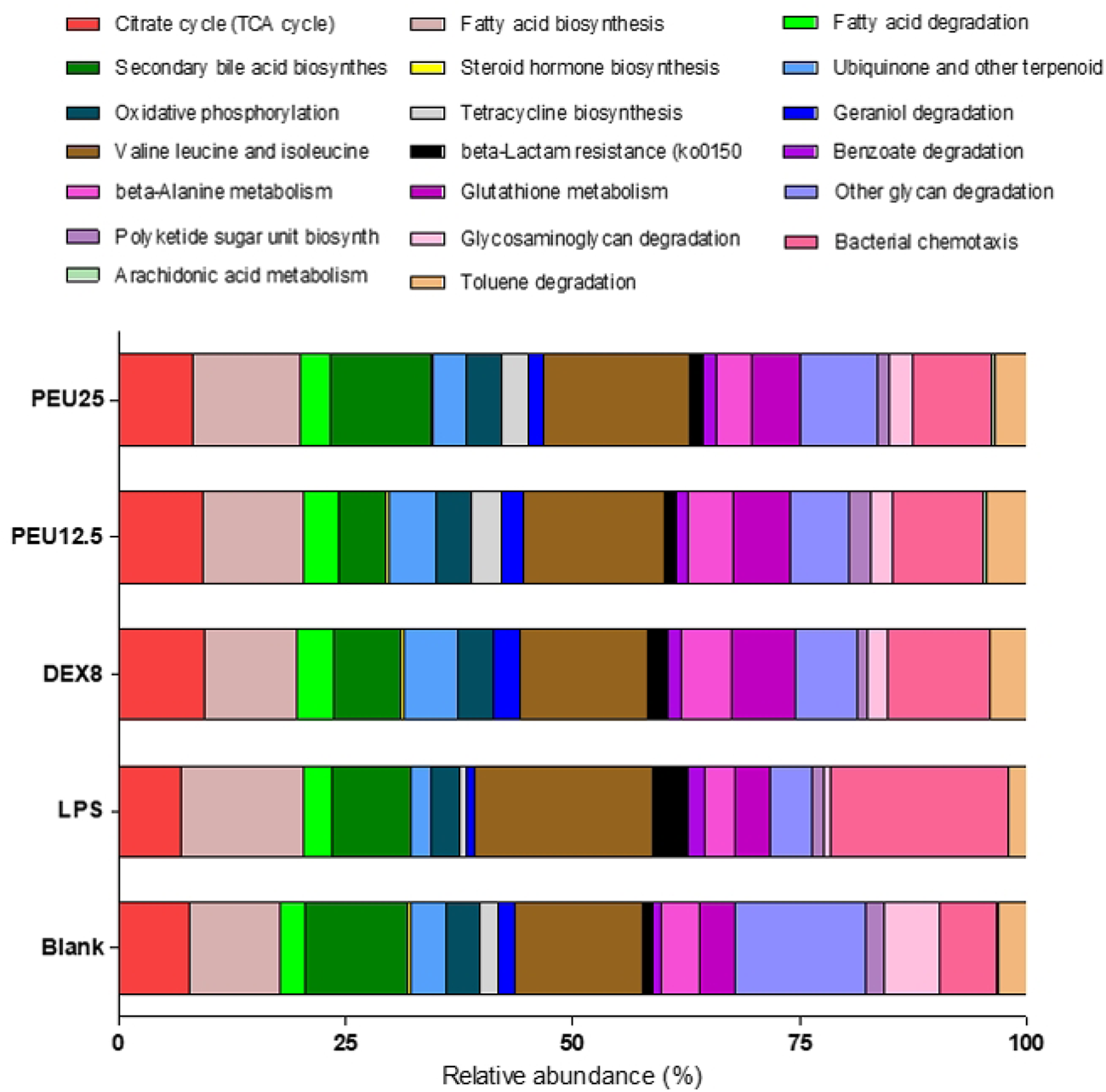
Flora composition at phylum and genus levels, and metabolism pathways in intestinal flora. (x̅ ±s, n=5-8). \**p*<0.05, \*\**p*<0.01 vs. Blank; ^#^*p*<0.05, ^##^*p*<0.01 vs. LPS. **A,** Changes in flora composition at phylum level. The significantly changed flora were shown to reflect their proportions in the community at the phylum level; **B**, Changes in flora composition at genus level. The proportions of significantly altered flora were shown in the community at the genus level; **C**, the significantly changed metabolism pathways were shown in intestinal flora at early stage of sepsis after treatments.

At the genus level, 285 bacteria were markedly altered after the treatments (*p* < 0.05). The PEU treatment significantly increased the proportions of *Nesterenkonia* and *Rikenellaceae*_RC9_gut_group, and decreased the proportions of *Vibrio*, *Pseudoalteromonas*, *Colidextribacter*, *Flaviflexus*, *Faecalibaculum*, *Enterorhabdus*, and *Ralstonia*, *Lachnospiraceae*_NK4B4_group compared to the LPS treatment at the genus level (Figure 7B). It suggested that the PEU treatment promoted some beneficial bacterial growth and inhibited some harmful bacterial growth, which to some extent improved disordered composition of the intestinal flora during sepsis.

In addition, 125 metabolism pathways were remarkably affected after the treatments (*p* < 0.05). The metabolism pathway analysis showed that PEU significantly affected energy metabolism, fatty acid degradation, secondary bile acid biosynthesis, other glycan degradation, toluene degradation, beta-lactam resistance, bacterial and chemotaxis in the intestinal flora at the early stage of sepsis (Figure 7C).

## Discussion

Up to now, few studies have reported the anti-inflammatory effect and the underlying mechanisms of PEU. To clarify targeting properties of PEU to the PAMPs, the ITC method is used to measure the binding affinities of this compound to KLA, ODN1826, PGN that are vital to initiate sepsis.

Currently, LPS (Lipid A) [17], and CpG DNA [18, 19] are used to test the anti-sepsis activity of extracts or compounds isolated from TCMs by biosensor technology. Also, anti-PGN antibodies [20] and complement C5 [21] have been applied to treat Gram-positive bacteria-induced sepsis.

The ITC is an important method to study bio-thermodynamics and biokinetics. It can continuously and accurately monitor and record energy changes through a highly sensitive automatic microcalorimeter, which provides in situ, online thermodynamic, and kinetic information simultaneously. Recently, the ITC method has been widely applied to measure binding of protein-protein, protein-DNA, or protein-small molecules [22–24].

PEU had moderate binding affinities to KLA and ODN1826 at a μM level but not PGN, suggesting its therapeutical potential in Gram-negative-induced infection. LPS activates membranous TLR4, thereby leading to activation of MyD88 and then NF-κB, which results in releases of proinflammatory cytokines crucial to the development of sepsis. Similarly, when free CpG DNAs are captured by cellular endocytosis, local plasma membrane collapse forms alveole and then vesicle. The vesicles containing CpG DNAs then incorporate endosomes to form larger endosomes where CpG DNA binds to TLR9 to activate the TLR9/MyD88/NF-κB signal pathway as well as TLR4, producing large number of proinflammatory cytokines [25]. Here, the ITC assay preliminarily confirmed the targeting properties of PEU to these two PAMPs.

*In vitro,* PEU inhibited the TLR4/MyD88/NF-κB pathway and reduced the down-stream proinflammatory cytokines TNF-α and IL-6 levels. *In vivo*, PEU protected CLP-induced sepsis mice, and reduced the serum TNF-α and IL-6 levels. Importantly, PEU attenuates the pathological injuries of the intestines. Further, PEU significantly upregulated the CLP-induced decreased intestine proliferation-related proteins Bmil1 and Lgr5.

The Bmi1 gene belongs to the Polycomb group family, and it is one of the members of Poly comb representational complex 1 (PRC1). It plays an important role in morphogenesis and hematopoietic processes during embryonic development [26]. Bmi1+ cells are reserve stem cells (RSCs) for small intestinal epithelial injury, and they can replace and supplement leukemia rich repeat containing G protein coupled receptor 5+ (Lgr5+) deficient intestinal stem cells (ISCs), thereby avoiding depletion of active ISCs in the circulation and preventing accumulation of damaged cells which accelerate tumor development [26]. At present, a study has reported that Bmi1 maintains intestinal tight junctions, epithelial barrier function, and microbial balance by preventing aging characterized by the accumulation of cell cycle inhibitory factor P16INK4a [27].

Lgr5 belongs to a rhodopsin-like family of G-protein-coupled receptors (GPCRs), and it is expressed on the cell surface and exists in various tissues such as the intestine, hair follicles, and skin. The Lgr5 contributes greatly to the steady-state regeneration of small intestine and colon. Compared to colon, the more active division of the Lgr5+ cells in small intestine reflects difference in the transformation rate of the two organ epithelial cells [28]. In addition, the self-renewal and high differentiation potential of Lgr5+ ISCs play an important role in the repairing of intestinal mucosal injury [29].

Until now, the anti-inflammatory effect of PEU and the role of TLR4/MyD88/NF-κB pathway in this event has not been clarified yet. To confirm the correlation between this compound and the pathway, chloroquine (CQ), a specific blocker of TLRs was used to validate the underlying mechanism. CQ is an antimalarial and anti-inflammatory agent, which is widely used to treat malaria [30, 31] and rheumatoid arthritis [32]. In addition, it is TLR3, 4, 8, and 9 inhibitors [33–36] and an autophagy antagonist [37, 38]. CQ reduced mortality and acute kidney injury of a polymicrobial sepsis mouse model by regulating serum cytokines TNF-α and IL-10 [36]. Moreover, CQ inhibits the TLR4/MyD88 signaling pathway in LPS-induced RAW264.7 cells [35].

Our findings showed that *in vitro* the PEU plus CQ further reduced the supernatant TNF-α and IL-6 levels significantly compared to PEU or CQ alone. *In vivo*, it showed that PEU remarkably protected the CLP-induced sepsis mice. However, there were no significant in the survival rate among the PEU, CQ, and CQ plus PEU groups. Moreover, the CQ plus PEU further significantly reduced the serum TNF-α and IL-6 compared to the CQ or PEU alone. These results indicated that the anti-inflammatory effects of PEU were further enhanced *in vitro* and *in vivo* in the presence of CQ, suggesting an independent role of the TLR4/MyD88/NF-κB pathway and involvements of other signal pathways in this anti-inflammatory event of PEU. In addition, the PEU plus CQ didn’t further elevate the survival of the sepsis mice compared to the PEU or CQ alone although the combined treatment further significantly reduced the serum cytokines, suggesting some irrelevance of cytokine levels and survival rate.

Currently, it has shown that the occurrence and development of sepsis was closely correlated to the intestinal flora imbalance [39]. The imbalance during sepsis will result in changes in composition, activity, and distribution of the flora, which induces inflammatory and oxidative stress in upper intestine, abnormal intestinal immune function, intestinal microcirculation disorders, and decreases products of short chain fatty acids (SCFAs) [40]. Further, the severity of sepsis is dependent on degree of the intestinal flora imbalance [41].

The present study revealed no significant differences in α diversity-related indices including Chao1, Shannon, and Simpson between the blank group and the LPS-treated group, which was basically consistent with a study on the intestinal flora of LPS-induced sepsis at the early stage [42]. However, our finding was opposite to the intestinal flora in a CLP-induced sepsis mouse model from 6 h to 3 days [43]. In this study, it showed that the CLP surgery significantly decreased the Chao1, Ace, Shannon indices, and increased the Simpson index compared to the sham along with the prolongation of modeling time, suggesting decreased richness and diversity of the intestinal flora after the CLP surgery. Here, we speculated that the differences might be due to the modeling methods and duration of the observation time.

In the present study, PEU significantly increased the Chao1 but not the Shannon and Simpson indices, which suggested that PEU in some degree improved species richness of the intestinal flora at the early stage of sepsis. Further, PEU increased the proportions of some beneficial bacteria and reduced the proportions of some harmful bacteria at the phylum and genus levels. Further, the PEU treatment significantly changed some metabolism pathways. For example, PEU enhanced TCA cycle inhibited adhesive invasive bacteria and reduced their virulence, increased lipoic acid metabolism to accelerate clearance of toxic metabolites, raised secondary bile acid biosynthesis to increase the SCFAs’ products, elevated oxidative phosphorylation level to provide energy for the body, decreased bacterial chemotaxis and antimicrobial drug resistance to reduce the bacterial motility, and decreased peptidoglycan biosynthesis to eliminate the toxins.

In summary, PEU has moderate binding affinities to KLA and ODN 1826, and exerts anti-inflammatory effects in sepsis-related models independent of the TLR4/MyD88/NF-κB pathway *in vitro* and *in vivo*. Moreover, PEU improves the disordered intestinal flora imbalance in the LPS-induced sepsis mouse model in some degree at the early stage of sepsis. This study will provide some evidences for the application of this compound in the treatment of sepsis-induced intestinal mucosal injury and dysbiosis.

## Funding

This work was supported by grants from 2022 High-Level Talent Research Project of Yunnan Provincial Health Commission (2023-KHRCBZ-A02), and Open Projects for Construction Unit of Clinical Pharmacy Center of Yunnan Province (2023YJZX-YX11).

## Acknowledgements

The authors would like to thank Fengzi Biotechnology Co., Ltd (Nanjing, China) for its technical support in 16S rRNA sequencing.

## Competing interests

All the authors declared no conflict of interest in this study.

## Author contributions

Qi Yao designed this study; Qi Yao did part of western blotting assay, ELISA assays *in vitro* and *in vivo*, animal survival assay, and wrote this manuscript; Bo-tao Chang performed part of western blotting assay and did the pathological examination.

## Data Availability Statement

The data in this study are deposited in the figshare repository, and the accession number is 10.6084/m9.figshare.26363479.

## Notes

### Competing Interest Statement

The authors have declared no competing interest.

## References

[1] Fleischmann C, Scherag A, Adhikari NK, Hartog CS, Tsaganos T, Schlattmann P, Angus DC, Reinhart K; International Forum of Acute Care Trialists. Assessment of global incidence and mortality of hospital-treated sepsis. Current estimates and limitations. Am J Respir Crit Care Med. 2016; 193:259–272.

[2] Labib A. Sepsis Care Pathway 2019. Qatar Med J 2019: 4.

[3] Wiens MO, Kissoon N, Holsti L. Challenges in pediatric post-sepsis care in resource limited settings: a narrative review. Transl Pediatr. 2021; 10:2666–2677.

[4] Musie E, Moore CC, Martin EN, Scheld WM. Toll-like receptor 4 stimulation before or after Streptococcus pneumoniae induced sepsis improves survival and is dependent on T-cells. PLoS One. 2014; 9: e86015.

[5] Kumar V. Toll-like receptors in sepsis-associated cytokine storm and their endogenous negative regulators as future immunomodulatory targets. Int Immunopharmacol. 2020; 89:107087.

[6] Schimmer RC, Urner M, Voigtsberger S, Booy C, Roth Z’Graggen B, Beck-Schimmer B, Schläpfer M. Inflammatory kidney and liver tissue response to different hydroxyethylstarch (HES) preparations in a rat model of early sepsis. PLoS One. 2016; 11: e0151903.

[7] Hu Q, Ren H, Li G, Wang D, Zhou Q, Wu J, Zheng J, Huang J, Slade DA, Wu X, Ren J. STING-mediated intestinal barrier dysfunction contributes to lethal sepsis. EBioMedicine. 2019; 41: 497–508.

[8] Graspeuntner S, Waschina S, Künzel S, Twisselmann N, Rausch TK, Cloppenborg-Schmidt K, Zimmermann J, Viemann D, Herting E, Göpel W, Baines JF, Kaleta C, Rupp J, Härtel C, Pagel J. Gut Dysbiosis with bacilli dominance and accumulation of fermentation products precedes late-onset sepsis in preterm infants. Clin Infect Dis. 2019; 69: 268–277.

[9] Wang H, Wang Q, Chen J, Chen C. Association among the gut microbiome, the serum metabolomic profile and RNA m6A methylation in sepsis-associated encephalopathy. Front Genet. 2022; 13: 859727.

[10] Stadlbauer V, Horvath A, Komarova I, Schmerboeck B, Feldbacher N, Klymiuk I, Durdevic M, Rainer F, Blesl A, Stiegler P, Leber B. Dysbiosis in early sepsis can be modulated by a multispecies probiotic: a randomised controlled pilot trial. Benef Microbes. 2019; 10: 265–278.

[11] Kim SM, DeFazio JR, Hyoju SK, Sangani K, Keskey R, Krezalek MA, Khodarev NN, Sangwan N, Christley S, Harris KG, Malik A, Zaborin A, Bouziat R, Ranoa DR, Wiegerinck M, Ernest JD, Shakhsheer BA, Fleming ID, Weichselbaum RR, Antonopoulos DA, Gilbert JA, Barreiro LB, Zaborina O, Jabri B, Alverdy JC. Fecal microbiota transplant rescues mice from human pathogen mediated sepsis by restoring systemic immunity. Nat Commun. 2020; 11: 2354.

[12] Macareño-Castro J, Solano-Salazar A, Dong LT, Mohiuddin, M, Espinoza JL. Fecal microbiota transplantation for Carbapenem-Resistant Enterobacteriaceae: A systematic review. J Infect. 2022; 84: 749–759.

[13] Hong MJ, Kim J. Determination of the absolute configuration of khellactone esters from Peucedanum japonicum roots. J Nat Prod. 2017; 80: 1354–1360.

[14] Yao Q, Gao Y, Lai C, Wu C, Zhao CL, Wu JL, Tang DX. The phytochemistry, pharmacology and applications of Melicope pteleifolia: A review. J Ethnopharmacol. 2020; 251: 112546.

[15] Zhang C, Li Y, Yin C, Zheng J, Liu G. In vitro study on the effect of peucedanol on the activity of cytochrome P450 enzymes. Pharm Biol. 2021; 59: 935–940.

[16] Karunai Raj M, Balachandran C, Duraipandiyan V, Agastian P, Ignacimuthu S. Antimicrobial activity of Ulopterol isolated from Toddalia asiatica (L.) Lam.: a traditional medicinal plant. J Ethnopharmacol. 2012; 140:161–165.

[17] Chen GR, Zhang G, Li MY, Jing J, Wang J, Zhang X, Mackie B, Dou DQ. The effective components of Huanglian Jiedu Decoction against sepsis evaluated by a lipid A-based affinity biosensor. J Ethnopharmacol. 2016; 186: 369–376.

[18] Liu X, Cheng J, Zheng X, Chen Y, Wu C, Li B, Fu J, Cao H, Lu Y, Li J, Zheng J, Zhou H. Targeting CpG DNA to screen and isolate anti-sepsis fraction and monomers from traditional Chinese herbs using affinity biosensor technology. Int Immunopharmacol. 2009; 9:1021–1031.

[19] Liu X, Zheng X, Wang N, Cao H, Lu Y, Long Y, Zhao K, Zhou H, Zheng J. Kukoamine B, a novel dual inhibitor of LPS and CpG DNA, is a potential candidate for sepsis treatment. Br J Pharmacol. 2011; 162:1274–1290.

[20] Sun D, Raisley B, Langer M, Iyer JK, Vedham V, Ballard JL, James JA, Metcalf J, Coggeshall KM. Anti-peptidoglycan antibodies and Fcγ receptors are the key mediators of inflammation in Gram-positive sepsis. J Immunol. 2012; 189: 2423–2431.

[21] Keshari RS, Popescu NI, Silasi R, Regmi G, Lupu C, Simmons JH, Ricardo A, Coggeshall KM, Lupu F. Complement C5 inhibition protects against hemolytic anemia and acute kidney injury in anthrax peptidoglycan-induced sepsis in baboons. Proc Natl Acad Sci U S A. 2021; 118: e2104347118.

[22] Calamini B, Ferry G, Boutin JA. Melatonin binding to human NQO2 by isothermal titration calorimetry. Methods Mol Biol. 2022; 2550: 305–314.

[23] Reinstein O, Yoo M, Han C, Palmo T, Beckham SA, Wilce MC, Johnson PE. Quinine binding by the cocaine-binding aptamer. Thermodynamic and hydrodynamic analysis of high-affinity binding of an off-target ligand. Biochem. 2013; 52: 8652–8662.

[24] Slavkovic S, Johnson PE. Analysis of aptamer-small molecule binding interactions using isothermal titration calorimetry. Methods Mol Biol. 2023; 2570: 105–118.

[25] Peter ME, Kubarenko AV, Weber AN, Dalpke AH. Identification of an N-terminal recognition site in TLR9 that contributes to CpG-DNA-mediated receptor activation. J Immunol. 2009; 182: 7690–7697.

[26] Yan KS, Chia LA, Li X, Ootani A, Su J, Lee JY, Su N, Luo Y, Heilshorn SC, Amieva MR, Sangiorgi E, Capecchi MR, Kuo CJ. The intestinal stem cell markers Bmi1 and Lgr5 identify two functionally distinct populations. Proc Natl Acad Sci U S A. 2012; 109(2): 466–471.

[27] Wang Y, Zang X, Wang Y, Chen P. High expression of p16INK4a and low expression of Bmi1 are associated with endothelial cellular senescence in the human cornea. Mol Vis. 2012; 18: 803–815.

[28] Salehzadeh S, Hasanzad M, Kadijani AA, Akbari A. The Expression Analysis of Intestinal Cancer Stem Cell Marker Lgr5 in Colorectal Cancer Patients and the Correlation with Histopathological Markers. J Gastrointest Cancer. 2020; 51(2): 591–599.

[29] Zheng L, Duan SL, Wen XL, Dai YC. Molecular regulation after mucosal injury and regeneration in ulcerative colitis. Front Mol Biosci. 2022; 9: 996057.

[30] Boechat N, Carvalho RCC, Ferreira MLG, Coutinho JP, Sa PM, Seito LN, Rosas EC, Krettli AU, Bastos MM, Pinheiro LCS. Antimalarial and anti-inflammatory activities of new chloroquine and primaquine hybrids: Targeting the blockade of malaria parasite transmission. Bioorg Med Chem. 2020; 28:115832.

[31] Gao P, Liu YQ, Xiao W, Xia F, Chen JY, Gu LW, Yang F, Zheng LH, Zhang JZ, Zhang Q, Li ZJ, Meng YQ, Zhu YP, Tang H, Shi QL, Guo QY, Zhang Y, Xu CC, Dai LY, Wang JG. Identification of antimalarial targets of chloroquine by a combined deconvolution strategy of ABPP and MS-CETSA. Mil Med Res. 2022; 9: 30.

[32] Trenkic Božinovic MS, Stankovic Babic G, Petrovic M, Karadžic J, Šarenac Vulovic T, Trenkic M. Role of optical coherence tomography in the early detection of macular thinning in rheumatoid arthritis patients with chloroquine retinopathy. J Res Med Sci. 2019; 24: 55.

[33] Cui G, Ye X, Zuo T, Zhao H, Zhao Q, Chen W, Hua F. Chloroquine pretreatment inhibits toll-like receptor 3 signaling after stroke. Neurosci Lett. 2013; 548: 101–104.

[34] Hu J, Wang X, Chen X, Fang Y, Chen K, Peng W, Wang Z, Guo K, Tan X, Liang F, Lin L, Xiong Y. Hydroxychloroquine attenuates neuroinflammation following traumatic brain injury by regulating the TLR4/NF-κB signaling pathway. J Neuroinflammation. 2022; 19: 71.

[35] Wang YY, Liu X, Cao, HW, Wang N, Lu YL, Zheng J. Inhibition of chloroquine on TLR4-MyD88-independent signaling pathway in lipopolysaccharide/endotoxin-induced RAW264.7 cells. ACTA ACADEMIAE MEDICINAE MILITARIS TERTIAE. 2010; 32: 869–872.

[36] Yasuda H, Leelahavanichkul A, Tsunoda S, Dear JW, Takahashi Y, Ito S, Hu X, Zhou H, Doi K, Childs R, Klinman DM, Yuen PS, Star RA. Chloroquine and inhibition of Toll-like receptor 9 protect from sepsis-induced acute kidney injury. Am J Physiol Renal Physiol. 2008; 294: F1050–F1058.

[37] Xu J, Yang KC, Go NE, Colborne S, Ho CJ, Hosseini-Beheshti E, Lystad AH, Simonsen A, Guns ET, Morin GB, Gorski SM. Chloroquine treatment induces secretion of autophagy-related proteins and inclusion of Atg8-family proteins in distinct extracellular vesicle populations. Autophagy. 2022; 18: 2547–2560.

[38] Cocco S, Leone A, Roca MS, Lombardi R, Piezzo M, Caputo R, Ciardiello C, Costantini S, Bruzzese F, Sisalli MJ, Budillon A, De Laurentiis M. Inhibition of autophagy by chloroquine prevents resistance to PI3K/AKT inhibitors and potentiates their antitumor effect in combination with paclitaxel in triple negative breast cancer models. J Transl Med. 2022; 20:290.

[39] Wetter LA, Hamadeh RM, Griffiss JM, Oesterle A, Aagaard B, Way LW. Differences in outer membrane characteristics between gallstone-associated bacteria and normal bacterial flora. Lancet. 1994; 343: 444–448.

[40] Yang XJ, Liu D, Ren HY, Zhang XY, Zhang J, Yang XJ. Effects of sepsis and its treatment measures on intestinal flora structure in critical care patients. World J Gastroenterol. 2021; 27: 2376–2393.

[41] Cabrera-Perez J, Badovinac VP, Griffith TS. Enteric immunity, the gut microbiome, and sepsis: Rethinking the germ theory of disease. Exp Biol Med (Maywood). 2017; 242: 127–139.

[42] Li W, Lin MR, Guo XY, Chen HY, Wen D, Lin JD. Effect of rhubarb on early intestinal flora in septic mice. J Snake (Science & Nature). 2021; 33: 138–140.

[43] Zhao H, Kuang C, Li F, Leng Y. Composition and changes of intestinal flora in septic mouse model. Zhonghua Wei Zhong Bing Ji Jiu Yi Xue. 2021; 33: 10–16.

